# Widely circulating pyrethroid resistance mechanisms reduce the efficacy of transfluthrin and pose a risk for mosquito-borne disease control with spatial emanators

**DOI:** 10.64898/2026.03.16.712035

**Authors:** Emmanouil Kokkas, Jeff Jones, David Weetman, Gareth Lycett, Mark J.I. Paine, Eirini Anastasaki, Walter Fabricio Martins, Timothy Hill, Ruth Cowlishaw, Amalia Anthousi, Fraser Colman, Hanafy Ismail, Rhiannon A.E. Logan, Panagiotis Milonas, John Vontas, Hilary Ranson, Rosemary S. Lees, Linda Grigoraki

## Abstract

Spatial emanators (SE) are a promising complement to existing tools for preventing mosquito transmitted diseases. In 2025, the WHO updated the WHO Guidelines for Malaria to include a conditional recommendation for the indoor use of prequalified SE products in malaria control. Both prequalified, and many other SE products contain the volatile pyrethroid transfluthrin, which shares the same target site as other (contact/solid phase) pyrethroids. Therefore, an assessment of cross resistance is critical to predict effectiveness against mosquitoes with existing pyrethroid resistance. Our results show that resistance to solid phase pyrethroids is correlated with resistance to transfluthrin in *Anopheles* and *Aedes* species. Moreover, commonly-selected resistance mechanisms including target site mutations and over-expression of P450 detoxification enzymes can confer resistance to transfluthrin. Furthermore, we show that resistant mosquitoes are less impacted by transfluthrin in terms of flight activation (irritancy) and reduced blood feeding inhibition, with the response correlating with resistance strength. Transfluthrin did not elicit an electroantennography response in *Anopheles gambiae* and surgically ablating mosquitoes’ antennae did not result in differences in flight activation upon transfluthrin exposure, suggesting the antennae are not required for transfluthrin to elicit behavioral responses. These results provide new insight regarding the mode of action of transfluthrin and the risk of resistance reducing transfluthrin’s efficacy in vector control interventions.

## Introduction

Vector borne diseases (VBD) are a growing public health problem in many parts of the world, with developing countries carrying the heaviest burden. Malaria cases remain at unacceptably high levels, with 597,000 deaths reported in 2023^1^, while dengue and other arboviral diseases are on the rise^2^. Vector control interventions that rely on contact insecticides to target indoor feeding and resting mosquitoes, including insecticide treated bed nets (ITNs) and indoor residual spraying (IRS), have contributed significantly to the reduction of VBD^3^. However, these interventions are often logistically difficult to implement at a national scale and their efficiency is threatened by the growing prevalence and intensity of resistance to contact insecticides^4^. Furthermore, there is a major gap in protection from outdoor transmission. Even in the case of malaria, transmitted by endophagic *Anopheles* mosquitoes, it is now well established that outdoor biting contributes to the transmission of the disease^5^. Thus, it is widely acknowledged that additional tools are needed to successfully combat VBD.

Spatial emanators (SEs), also known as spatial repellents (SRs), have been proposed as effective tools to address protection gaps in both indoor and outdoor settings^6,7^. They are widely used in the form of mosquito coils, vaporizers, emanators, or heated mats for personal protection^7,8^. SEs are compounds that act in the vapour phase disrupting the ability of mosquitoes to detect and feed on their hosts. Among the available SEs, transfluthrin has received most attention. The World Health Organization (WHO) has prequalified two similar transfluthrin-based spatial emanators for public health use^9^, with their large scale deployment expected in the next period. Transfluthrin is a volatile pyrethroid that, like all other members of this insecticide class, binds to voltage gated sodium channels (VGSCs) inhibiting their inactivation. This disrupts the generation of the membrane potential in nerve cells, impairing their normal function. At high concentrations, transfluthrin is neurotoxic causing paralysis and death. At lower concentrations, it likely causes neuronal hyperexcitation that is manifested by behavioral responses such as repellency and inhibition of blood feeding ^10–12^. However, the mechanistic basis of its repellent action remains poorly understood, including whether and how transfluthrin is detected by the mosquitoes’ sensory organs.

Transfluthrin-based interventions, in the form of active and passive emanators, treated sandals and hessian strips have been tested in a number of field studies. These studies demonstrated personal protection against mosquito vectors, evidenced by significant reductions in human-vector contact rates and/or mosquito density^13–15^. Likewise, a limited number of studies assessing the epidemiological impact of transfluthrin emanators, showed reduction in malaria^6,16,17^ and dengue^18^ cases. The promising outcomes of those studies have recently led WHO to issue a conditional recommendation supporting the use of spatial emanators for malaria control as a supplemental tool for use indoors^9^. However, to fully assess the efficacy of transfluthrin, further research is needed to evaluate its impact on disease transmission across diverse ecological settings. Crucially, it is also necessary to determine whether, and to what extent, cross resistance to widely used contact pyrethroids could compromise both the toxic and repellent effects of transfluthrin.

Mutations in the common VGSC target site, known as the knockdown resistance (kdr), have long been shown to cause pyrethroid resistance to contact insecticides. A limited number of published studies have shown an association between high or increasing frequencies of the 995F kdr mutation and reduced mortality^19,20^ as well as behavioral avoidance^20^ in response to transfluthrin. However, other studies found no association between target site resistance mutations and the efficacy of volatile pyrethroids^21,22^. Furthermore, insights from mutational analysis of *Aedes aegypti* sodium channels and computational modeling suggest that transfluthrin has a different binding site within the VGSC compared to contact pyrethroids^23^. Under this scenario, mutations conferring resistance to contact pyrethroids may not equally affect transfluthrin binding, limiting the extent of cross resistance.

It remains unclear whether transfluthrin is prone to metabolism by the detoxification enzymes commonly over-expressed in pyrethroid-resistant populations. One of the major routes of pyrethroid metabolism is oxidation of the phenoxybenzyl moiety common to members of this insecticide class^24,25^. Such sites are blocked by polyfluorination of the benzyl ring in transfluthrin, suggesting that typical P450-mediated metabolism, a key mechanism of pyrethroid resistance, may be hindered. This hypothesis is supported by studies showing that an *Anopheles funestus* strain, highly resistant to contact pyrethroids due to elevated P450 activity, remains susceptible to transfluthrin^26^. Additionally, P450 inhibitors, including piperonyl butoxide (PBO), have not been able to increase susceptibility to transfluthrin ^26,27^. However, several other routes of metabolism by P450s are feasible ^24,28^. Furthermore, over-expression of specific P450s such as CYP6M2 and CYP9K1, that are known to metabolise contact pyrethroids, has been observed following the selection of an *An. gambiae* strain with transfluthrin^19^. These findings suggest that while transfluthrin may evade some common resistance mechanisms, its interaction with metabolic pathways is complex and warrants further investigation.

Overall, the current literature indicates that mosquitoes have the capacity to evolve both physiological and behavioral resistance to transfluthrin. However, the association between resistance to contact pyrethroids and impaired transfluthrin efficacy remains ambiguous. In this study we use non-contact assays to quantify the insecticidal effect of transfluthrin in pyrethroid sensitive and resistant *Aedes* and *Anopheles* mosquitoes, and to assess how resistance to contact pyrethroids correlates with resistance to transfluthrin. We further use a suite of transgenic and genome-edited *An. gambiae* lines to functionally validate the role of target site mutations and over-expression of P450s in conferring transfluthrin resistance. Additionally, we test the ability of P450s to metabolise transfluthrin *in vitro*. To explore how resistance may influence transfluthrin-elicited repellency, we analyse the flight behavior of *An. gambiae* mosquitoes during transfluthrin exposure, comparing responses between resistant and susceptible strains. Furthermore, we investigate the involvement of the mosquitoes’ primary olfactory organ, the antennae, in the detection and behavioral response to transfluthrin.

## Materials and Methods

### Strains used

#### Field colonized

*For Aedes aegypti* the strains used were: 1) Cayman, which was colonized around 2009 from field collections in the Grand Cayman^29^, 2) Cayman XX selected from the Cayman strain in the laboratory using 3% permethrin for several generations^30^, 3) Jeddah, a Saudi Arabian strain colonized in 2015^31^ and 4) New Orleans, a standard susceptible *Ae. aegypti* strain colonized over 50 years ago^30^. For *Anopheles gambiae* the strains used were: the highly pyrethroid resistant VK7 2014 and Tiassalé 13 originating from Burkina Faso and Cote d’Ivoire and colonized in 2014 and 2011 respectively^32,33^ and the standard susceptible strain (Kisumu) colonized over 40 years ago. For *Anopheles funestus* one strain was used, FuMOZ^26^ that is highly resistant to pyrethroids and its resistance is mediated by detoxification enzymes only^34^.

#### Transgenic/Genome edited *An. gambiae* lines

Two CRISPR edited *An. gambiae* lines, the Kisumu-995F and Kisumu-402L, carrying target site mutations at the voltage gated sodium channel were used. The generation of the lines is described in Grigoraki et al., 2021^35^ and Williams et al., 2022^36^ respectively. Both strains were generated using Kisumu as the standard susceptible background and both are homozygous for the introduced mutation. Kisumu-995F shows 14.6 fold resistance to deltamethrin, while Kisumu-402L shows 5 fold resistance.

The CFP marked Gal4 driver line Ubi-A10 (expressing Gal4 under the poly-ubiquitin promoter), generated by Adolfi et al., 2018^37^ was used to drive over-expression of P450s in multiple tissues (detailed description provided in Adolfi et al., 2018)^37^. This line was crossed with the responder lines UAS-Cyp6P3 and UAS-Cyp6M2 (both generated by Adolfi et al., 2019^38^ using G3 as the standard susceptible genetic background). Progeny of the crosses over-expressing Cyp6M2 or Cyp6P3 were used for toxicity bioassays and compared to progeny of the cross between the driver line and G3.

#### Rearing conditions

All mosquito strains were maintained at 26°C ± 2°C, 80% relative humidity ± 10% and under a L12:D12 hour light:dark cycle with a 1 hour dawn and dusk. Larvae were fed ground fish food (Tetramin tropical flakes,Tetra, Blacksburg, VA, USA) and adults maintained on 10% sucrose solution fed *ad libitumin*.

### Toxicity bioassays “Deli-pot” assay

Insecticides, deltamethrin and transfluthrin, were diluted in acetone (concentrations for deltamethrin ranging from 0 to 1,000μg/ml and for transfluthrin ranging from 0 to 2,000μg/ml) and applied (total volume of 1 ml) by pipette to glass Petri dishes (radius 2.5cm, area 19.635cm^2^, SLS, Nottingham, UK). Negative controls included acetone alone. Prior to testing the acetone was left to evaporate by putting treated dishes on an orbital shaker for 15 min. A plastic container (‘deli-pot’, 2oz size), with a hole in the top sealed by parafilm was fitted over each Petri dish. For non-contact assays, the base of the pot was sealed by mesh, while in contact assays deli-pots were covered at the base by a removable, rather than fixed mesh, which was gently withdrawn when the pot was applied to the glass dish (Supplementary Figure 1). Ten to fifteen female mosquitoes were aspirated into each plastic container through the hole and exposure proceeded for 30 min in an incubator. At the end of the assay knockdown was recorded, and mosquitoes were aspirated (using a mechanical aspirator) into paper cups and held for 24 h at insectary conditions (with provision of 10% sucrose) to record final mortality. All mosquitoes used in these assays were 2-5 days old, except for the data on Table 3 where 2-9 day old mosquitoes were used.

### Peet Grady assay Set-up

The desired quantity of transfluthrin (0.25, 5 and 10 mg) (99,1% purity, LGC, CAS:<u>118712-89-3</u>) was weighed and dissolved in 3 mL of acetone. The resulting solution was used to impregnate grade 1 Whatman filter papers (Cytiva; 14.5 × 12 cm), which were subsequently placed in plastic boxes for 10 min to allow complete solvent evaporation. The treated filter papers were then mounted within a custom-designed 3D-printed scaffold and suspended at the center of the Peet Grady chamber using a plastic hook. Groups of approximately 25 female non blood fed *An. gambiae* mosquitoes, 2–5 days old, were introduced into each side of a dual-compartment cage. Cages were positioned adjacent to the windows of the Peet Grady chamber and secured in place with plastic hooks. The assay was conducted as previously described^39^, with the modification that cameras were employed to record mosquito movement and behavior, which was subsequently analysed. Upon assay completion, mosquitoes were transferred to 350ml paper holding cups provisioned with cotton pads soaked in 10% sucrose solution and maintained for 24 h. Mortality was then assessed.

## Data analysis

### Machine Learning Method to Estimate the Number and Location of Mosquitoes

A deep-learning neural network classifier based on that devised in Redmon et al, 2021^40^ was trained on a hand-labelled dataset of images sampled from the video recordings of the Peet Grady chamber, obtained at 30 frames per second with 1.1 × zoom using an action camera (The Xtreme I + 4 K, Campark). The training set comprised a single class (“mosquito”) and all training images had mosquito locations localised within bounding boxes, resolution permitting. This data set was further augmented by randomly processing the images to improve the robustness of the classifier. The augmentation process included horizontal and vertical flip, adjustment of brightness and contrast, gamma adjustment, RGB channel shifts, image rotation (+-5° limit), and Gaussian blur. The final dataset was 2,150 images, randomly allocated to training, validation and test groups with proportions (0.7, 0.2, 0.1) respectively. Training was performed for over 1,000 epochs on a Windows 10 computer using an Nvidia 4060 Cuda hardware accelerator running a custom training method using PyTorch^41^. The performance parameters of the final trained network are shown in Supplementary Table 2.

The trained network was then used to evaluate previously unseen experimental recordings at 1-minute time intervals and any detected mosquitoes, and their screen coordinates (centroid of the bounding box detections), were appended to log files for further analysis.

Log files were analysed with custom R scripts to collate the number of detections per sample and their screen locations. For twin-cage experiments we used an algorithm to determine which sides of the cage detected mosquitoes were located (and therefore which strain category) based on comparison of the mosquito coordinates and coordinates of the polygonal seam boundary running vertically down the centre of the cage. Detections were also assigned to the categories ‘floor’ or ‘upper’ based on whether their *y* coordinate was below the seam representing boundary of the cage floor (supplementary figure 6).

Scripts to assemble, collate and analyse the data from all recordings were written in R ^42^ using the R-Studio development environment ^43^ and plots were generated using the GGPlot2 package^44^.

### Optical Flow Method for to Estimate Movement Activity

Optical Flow is a concept originally devised to model the detection of relative movement in visual systems of insects and higher animals^45^. More recently it has been used as part of computational pipelines for detection and classification of moving objects ^46^.

Video recordings of activity in the PG chamber were analysed by a bespoke motion estimation software program developed at LSTM (available upon request). The software allows a user to load a video recording, define a global mask around the cage to exclude unwanted regions. Individual polygonal regions of interest (ROI) can be created by the user to represent different regions of the cage from which optical flow can be measured. For this study, twin-compartment mosquito cages were used to improve the workflow. Each mosquito cage had two vertically divided compartments where two different mosquito population could be inserted and kept separate. Each cage half had two regions defined: upper and lower. The upper regions were all regions of the half-cage except the floor and the lower region defined the floor of the half-cage.

Video frames were compared at 5s intervals using the Farneback method of computing dense optical flow^47^ by greyframe pixel intensity difference at t_x_ and t*_x_-5.* A low pass 2D Gaussian filter (k=3) was used to smooth each image frame before optical flow assessment. A regular 2D Vector field of points representing vector heading and vector magnitude was generated across the image for each timestep. The values in the vector field were aggregated by ROI previously selected by the user (supplementary figure 6) and movement vectors inside each ROI were thresholded for minimum and maximum signal range. Minimum thresholds were used to filter out extraneous video noise contamination (for example random movement of cage netting, or compression artifacts at netting seams). Maximum thresholds were used to clip large sudden temporary regions of signal contamination (for example, any sudden movements of the equipment).

The mean vector field magnitude and heading for each ROI was appended to a text log data structure at each 5s sample interval which were saved at the end of each video analysis run. The text log files were later aggregated and analysed using bespoke R scripts to provide information about the mean total amount of movement and orientation of movement within cage ROIs and the movement activity of mosquitoes over time, during the duration of the entire 1h experiment.

### Recombinant P450 production and transfluthrin *in vitro* metabolism assay

Recombinant *E. coli* membranes co-expressing the *An. gambiae* cytochrome P450 enzymes CYP6M2, CYP6P3, or *An.* funestus CYP6P9a together with the *An. gambiae* NADPH cytochrome P450 oxidoreductase (AgCPR) were supplied by Cypex Ltd, UK (Dundee, UK) and using pCWori + expression vector constructs supplied by Dr Mark Paine as described previously for CYPs 6M2, 6P3^48^, and CYP6P9a^49^ produced according to established protocols ^48,49^. The manufacturer verified P450 expression, content and quality by Carbon Monoxide (CO)-difference spectroscopy, and CPR content was quantified via cytochrome *c* reductase activity assay. Protein content was estimated by the Bradford method, and samples were stored in aliquots at − 80 °C until use.

Metabolism assays were performed in 200 µL reaction mixtures containing 5 µM transfluthrin, 0.1 µM of the recombinant enzyme, 0.1 M of potassium phosphate buffer (pH 7.4), and 0.5 mM NADPH (Melford, UK) in the presence or absence of cytochrome *b5* (0.8 µM) supplied by Cypex Ltd ^24^. Reactions were incubated at 30°C with shaking (1200 rpm) for 2hours, then terminated by adding an equal volume (200 µl) of acetonitrile. The mixtures were vortexed and shaken for an additional 10 min, followed by centrifugation at 20,000 × g for 20 min to remove protein debris. A 100 µL aliquot of the clarified supernatant was injected onto a 250 mm × 4.6 mm C18 reverse-phase column (Acclaim 120, Thermo Scientific) for HPLC analysis. Separation was carried out under isocratic conditions using a mobile phase of 80% acetonitrile and 20% water, at a flow rate of 1 mL min⁻¹ and column temperature of 23°C. Transfluthrin eluted at 8.2 min and was quantified by peak area integration using the OpenLAB Chromatography Data System. Assays were conducted in three independent biological replicates; each performed in technical triplicate. Unpaired two-tailed t-test with Welch’s correction comparing +NADPH reactions and ± cytochrome *b5* were used to assess statistically significant substrate depletion.

### Blood feeding

For blood feeding post-transfluthrin exposure, mosquitoes were transferred to 350ml paper cups immediately after completion of the Peet Grady assay. A Hemotek feeder (*Hemotek Membrane Feeding System*. Blackburn, UK: Hemotek Ltd.) containing human blood preheated for 5 min was placed on top of the cups for 7 min. Each feeder was used no more than twice. Following feeding, mosquitoes were frozen for 10 min, after which blood-fed individuals were counted. Mosquitoes unable to fly during the feeding period (presumably dead or knocked down) were excluded from the total count. Post-exposure blood feeding was conducted immediately after chamber fumigation, within one hour of assay completion. For the 24-hour post-exposure blood feeding, mosquitoes were maintained in cups with cotton wool soaked in 10% sucrose solution for 24 hours after the assay. The sucrose source was then removed for at least 1 hour prior to blood feeding, which was performed using the same procedure described above.

### EAG

Electroantennogram recordings were performed using two glass capillaries filled with 0.1 M KCl solution. A silver wire connected to the recording device was inserted into the back of each capillary. Female *An. gambiae* (2–5 days old) were cold-anesthetized on ice for ∼1 min, and heads were excised with a fine scalpel. Antennal tips were cut to improve signal conduction. The head was mounted at the end of one capillary via the neck, while one antenna was inserted into the recording electrode, completing the circuit.

Odor stimuli were delivered using the Syntech Stimulus Controller CS-55 (Syntec, Kirchzarten, Germany). The chemicals used were transfluthrin (99,11% purity, LGC) as well as 1-octen-3-ol (>95% purity, LGC) and citronellal (95% purity, Aldrich) as positive controls. All test compounds were dissolved in acetone (Pesticide Grade, 99.8% purity, Carlo Erba) and 10μL were applied to 1 × 2 cm filter paper placed inside a glass Pasteur pipette. Acetone was allowed to evaporate for 1 minute before inserting the filter paper inside the pipette. During stimulation, a 1 s odor pulse was directed through the pipette and onto the antenna. Continuous airflow was maintained otherwise.

Signals were recorded with the Syntech IDAC-2 EAG GC-EAD Signal Acquisition Controller (Syntec, Kirchzarten, Germany). A recovery period of 2 min was allowed between successive stimuli. Each preparation was used for up to 30 min post-dissection, after which antennal responses declined. Data were analyzed with GcEad 32software (Syntec, Kirchzarten, Germany).

### Antennae ablation

Two-to five-day-old mosquitoes were anesthetized with CO₂, and their antennae were excised at the base using micro-scissors. After dissection, mosquitoes were transferred to 350ml paper cups supplied with cotton wool soaked in 10% sucrose solution and allowed to recover overnight. Surviving individuals were exposed to transfluthrin the following day.

## Results

### Development of a bench top toxicity bioassay for volatile pyrethroids and assessment of transfluthrin resistance in field colonized *Anopheles* and *Aedes* strains

Given that transfluthrin is a volatile pyrethroid that acts mainly in the vapor phase, we developed a simple assay to measure its toxic effect, by adapting the previously described tarsal plate assay (described in Lees et al., 2019^50^), used for contact insecticides. The assay is composed of a glass petri dish coated with transfluthrin and sealed with a mesh. A plastic container (deli-pot), in which mosquitoes are introduced, is placed on top of the mesh (Supplementary Figure 1). The mesh hinders mosquitoes from coming in direct contact with the transfluthrin treated surface, thus being exposed solely to the vapor phase.

The non-contact bioassay was used to assess the susceptibility of *An. gambiae*, *An. funestus* and *Ae. aegypti* field colonized strains to transfluthrin through dose response curves. Details of the origin, resistance status and resistance mechanisms present in each strain are provided in Supplementary Table 1.

Results for transfluthrin volatile assays are shown in Table 1. The LC_50_ and LC_90_ values, corresponding to the concentration that causes 50% and 90% mortality respectively, are shown for each strain. The RR_50_ (Resistance Ratio _50_= LC_50_ of resistant strain/LC_50_ of susceptible strain) and RR_90_ (Resistance Ratio _90_= LC_90_ of resistant strain/LC_90_ of susceptible strain) were calculated to define the strength of resistance. For the *Aedes* strains, Cayman showed no resistance (RR_50_ 1.3-fold), Jeddah showed moderate resistance (RR_50_ 7.6-fold) and Cayman XX showed high resistance (RR_50_ 57.4-fold). For the *Anopheles* strains, FuMOZ showed low levels of resistance (RR_50_ of 4.2-fold), while the two *An. gambiae* strains, Tiassalé 13 and VK7 2014 showed high resistance, with RR_50_ of 213.2-fold and 75.3-fold respectively.

**Table 1.** Resistance levels to transfluthrin and deltamethrin in different field colonized *Anopheles* and *Aedes* strains. Tarsal plate assays; non-contact and contact for transfluthrin and contact for deltamethrin were used to measure the LC_50_ (concentration that results in 50% mortality in ppm) and LC_90_ (concentration that results in 90% mortality in ppm) for each insecticide. Values are in ppm. Results are from probit regression analysis (accompanied by Supplementary Figures 2,3 and 4). LCL, UCL are upper and lower 95% confidence limits. RR_50_ and RR_90_ = resistance ratio at the 50 and 90 thresholds respectively; resistance ratios in bold are significantly greater than 1 (i.e. significant vs genus-specific susceptible strain, New Orleans or Kisumu).

| Genus | Strain | Transfluthrin volatile |  |  |  | Deltamethrin contact |  |  |  | Transfluthrin contact +volatile |  |  |  |
| --- | --- | --- | --- | --- | --- | --- | --- | --- | --- | --- | --- | --- | --- |
|  |  | LC<sub>50</sub><br>(LCL-UCL) | LC<sub>90</sub><br>(LCL-UCL) | RR<sub>50</sub> | RR<sub>90</sub> | LC<sub>50</sub><br>(LCL-UCL) | LC<sub>90</sub><br>(LCL-UCL) | RR<sub>50</sub> | RR<sub>90</sub> | LC<sub>50</sub><br>(LCL-UCL) | LC<sub>90</sub><br>(LCL-UCL) | RR<sub>50</sub> | RR<sub>90</sub> |
| *Aedes* | New Orleans | 0.33<br>(0.25-0.43) | 0.84<br>(0.61-1.56) | 1 | 1 | 0.02<br>(0.02-0.03) | 0.09<br>(0.06-0.19) | 1 | 1 | 0.05<br>(0.03-0.07) | 0.13<br>(0.09-0.29) | 1 | 1 |
| *Aedes* | Cayman | 0.42<br>(0.21-0.66) | 4.41<br>(2.93-7.95) | 1.3 | 5.2 | 0.26<br>(0.1-0.49) | 7.26<br>(4.08-17.13) | 11 | 78.9 | 0.26<br>(0.16-0.4) | 2.32<br>(1.39-5.08) | 5.5 | 17.7 |
| *Aedes* | Cayman XX | 18.83<br>(14.48-24.17) | 128.77<br>(86.33-232.32) | 57.4 | 152.8 | 7.79<br>(4.34-11.34) | 40.1<br>(25.66-94.02) | 324.6 | 435.9 | 0.86<br>(0.61-1.22) | 5.84<br>(3.56-12.71) | 18 | 44.6 |
| *Aedes* | Jeddah | 2.5<br>(1.63-3.7) | 10.57<br>(6.36-30.19) | 7.6 | 12.5 | 0.73<br>(0.3-1.41) | 9 (4-45.76) | 30.5 | 97.8 | 0.56<br>(0.27-1.24) | 5.5<br>(2.11-55.87) | 11.6 | 42 |
| *Anopheles* | Kisumu | 0.31<br>(0.23-0.42) | 0.97<br>(0.66-1.85) | 1 | 1 | 0.36<br>(0.26-0.49) | 2.74<br>(1.65-6.23) | 1 | 1 | 0.1<br>(0.08-0.12) | 0.22<br>(0.18-0.31) | 1 | 1 |
| *Anopheles* | Tiassalé 13 | 65.89<br>(48.99-86.47) | 366.23<br>(249.13-650.11) | 213.2 | 379.5 | 97.02<br>(59.61-201.98) | 815.75<br>(324.87-10017.8) | 271 | 298 | - | - | - | - |
| *Anopheles* | VK7 2014 | 23.27<br>(8.42-43.63) | 592.95<br>(265.48-2888) | 75.3 | 614.5 | 106.13<br>(68.4-171.6) | 547.36<br>(303.7-1534.47) | 296.5 | 200 | 1.4<br>(0.97-1.93) | 3.42<br>(2.39-7.29) | 14.2 | 15.5 |
| *Anopheles* | FuMOZ | 1.29<br>(0.93-1.86) | 3.97<br>(2.53-10.3) | 4.2 | 4.1 | 13.44<br>(8.52-19.67) | 77.83<br>(45.65-221.77) | 37.5 | 28.4 | 0.42<br>(0.03-0.74) | 1.34<br>(0.76-29.07) | 4.3 | 6.1 |

**Table 3.**
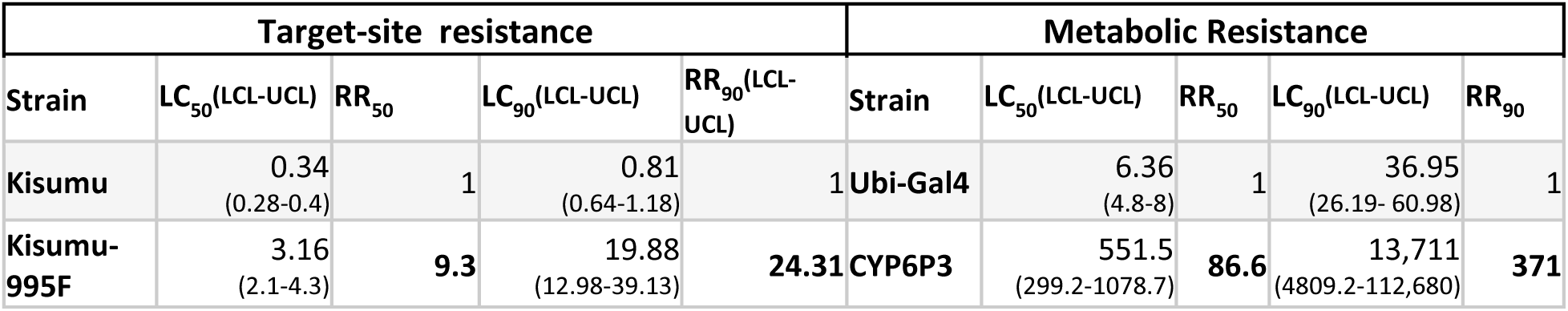
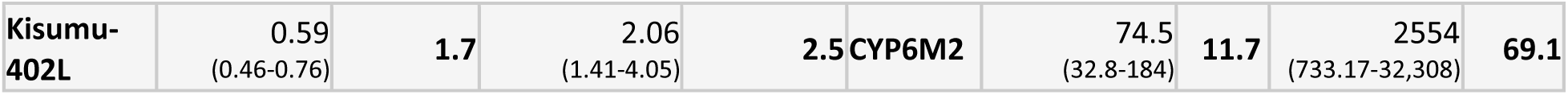
Resistance levels to transfluthrin conferred by single target site and metabolic resistance mechanisms. Non-contact tarsal plate assays to measure transfluthrin resistance in genome edited *Anopheles gambiae* lines carrying single VGSC point mutations and in transgenic strains over-expressing P450s. LC_50_ (concentration that results in 50% mortality) and LC_90_ (concentration that results in 90% mortality) values are given in ppm. Results are from probit regression analysis. LCL= 95% Lower confidence limit, UCL=95% Upper confidence limit. RR_50_ and RR_90_ = resistance ratio at the 50 and 90 thresholds respectively; resistance ratios in bold are significantly greater than 1 (i.e. significant vs susceptible strain, Kisumu and Ubi-Gal4).

To test if transfluthrin’s route of entry alters the resistance levels, we performed toxicity bioassays in which mosquitoes come in contact with transfluthrin’s vapour phase, as well as directly with the treated tarsal plate (same assay as before without the mesh). Resistance levels dropped for the Cayman XX and VK7 strains, increased slightly for Cayman and Jedah and remained unchanged for FuMOZ (Table 1).

### Resistance to the contact pyrethroid deltamethrin is correlated with resistance to transfluthrin

To assess how resistance to transfluthrin correlates quantitatively and qualitatively with resistance to contact pyrethroids, dose response assays to deltamethrin were performed for all strains, using the tarsal plate assay (without the mesh). Results for the tarsal contact assay with deltamethrin are shown in Table 1. All strains exhibited significant resistance compared to the reference strains, with RR_50_ resistance ratios greater than 10-fold. Amongst *Aedes* strains the permethrin-selected Cayman XX strain was far more resistant (RR_50_ of 324-fold) than its progenitor strain (Cayman) (RR_50_ of 11-fold), and also compared to the Jeddah strain which carries different target site resistance mutations (RR_50_ of 30.5-fold). The *An. funestus* FuMOZ strain showed substantial resistance (RR_50_ of 37-fold), while the two *An. gambiae* Tiassalé 13 and VK7 2014 strains showed very strong resistance with RR_50_ 271-fold and 296-fold respectively. Whilst RR estimates for the more moderately resistant strains Jeddah and FuMOZ were quite consistent at the 50% and 90% kill thresholds, that for Cayman was discordant, suggestive of a population with a mixed resistance profile.

### Assessing the levels of transfluthrin resistance conferred by single resistance mechanisms

To assess how transfluthrin toxicity is affected by the presence of two VGSC mutations that have been shown to confer resistance to contact pyrethroids^35,36^, we used two CRISPR edited *An. gambiae* strains that carry in a fully susceptible (Kisumu) genetic background either the 995F or V402L mutation in homozygosity. Homozygotes for 995F showed 9.3-fold resistance to transfluthrin, while homozygotes for V402L showed a much lower 1.7-fold resistance (Table 2). Furthermore, a Gal4 driver line^37^, that expresses the transcription activator Gal4 under the control of the polyubiquitin promoter (Ubi-Gal4) was used to drive over-expression of two P450 genes, *Cyp6M2* and *Cyp6P3*, upon crossing with the respective UAS lines^38^. Multi-tissue over expression of Cyp6P3 resulted in 86.6-fold resistance to transfluthrin and multi-tissue over-expression of Cyp6M2 in 11.7-fold resistance.

### Recombinantly expressed *Anopheles* P450s metabolize transfluthrin

The *An. gambiae* P450s, CYP6M2 and CYP6P3 and the *An. funestus* CYP6P9a were recombinantly expressed in *E. coli* together with *An. gambiae* NADPH P450 reductase (CPR). The ability of the expressed P450s to metabolise transfluthrin was evaluated by measuring insecticide turnover (% substrate depletion after 120 min of incubation) in the presence and absence of NADPH. Cytochrome b5 is also known to increase catalytic activity. Therefore, reactions were carried out in the presence and absence of b5 to determine b5 effects. A cut-off value of 20% substrate depletion was used to distinguish true substrate turnover from baseline variability ^49,51^. In the absence of b5 CYP6M2 showed a 57.16 ± 0.67 % depletion, while CYP6P3 and CYP6P9a showed less than 20%. However, the addition of b5 significantly enhanced transfluthrin metabolism for all three P450s. CYP6M2 showed 88.13 ± 4.48% depletion, CYP6P3 75.87 ± 6.75% and CYP6P9a 65.18 ± 1.53% (Figure 1). Representative HPLC-chromatograms of the transfluthrin metabolism assays are shown in Supplementary Figure 5. In addition to the depletion of the transfluthrin peak, two clear metabolite peaks with retention times at 5 min (M1) and 6 min (M2) were produced (Supplementary Figure 5).

**Figure 1.**
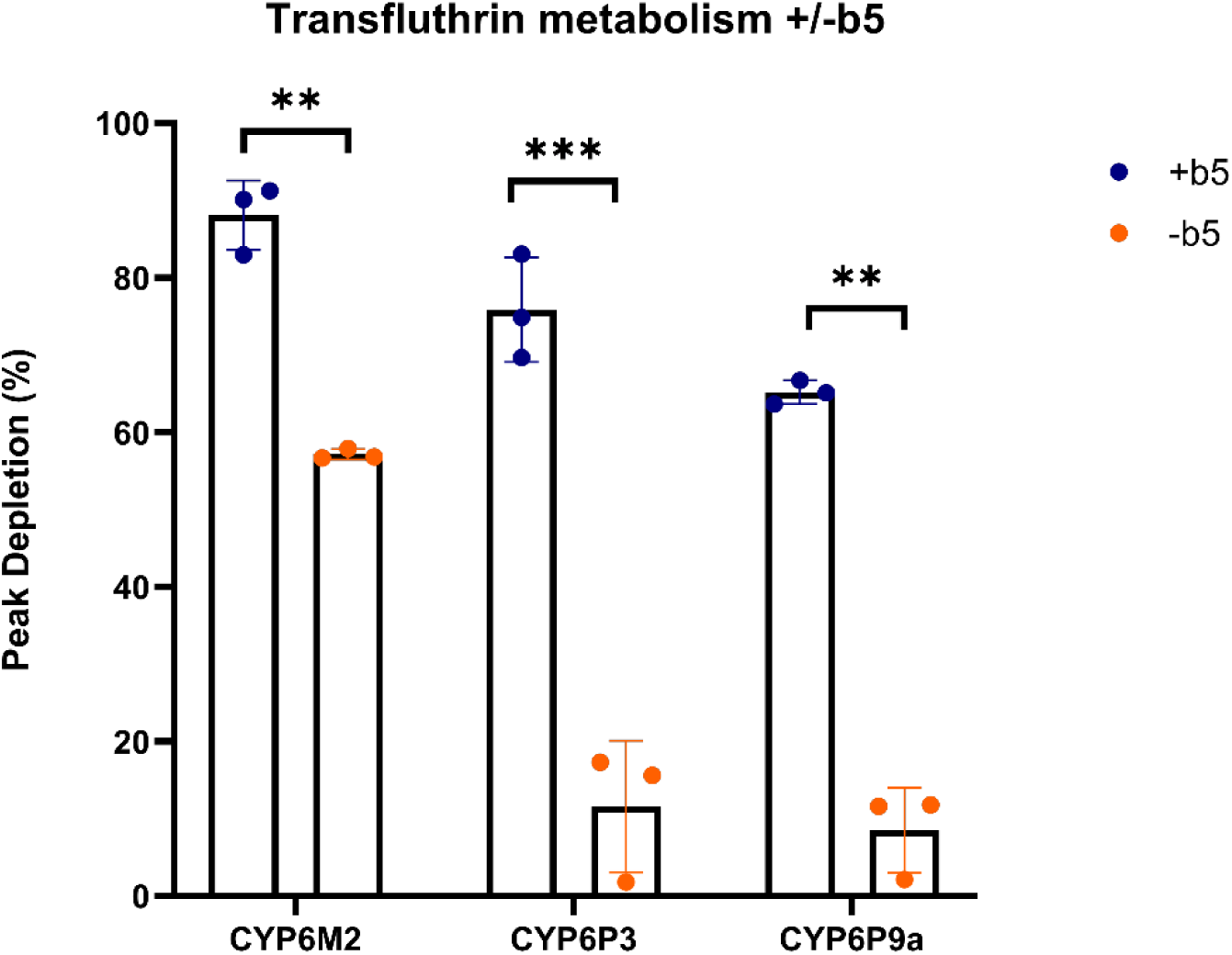
In vitro metabolism of transfluthrin by recombinantly expressed Anopheles P450s. Bars represent the proportion (% depletion) of 5μM insecticide cleared by 0.1μM P450 with (0.8μΜ) or without b5 in the presence of NADPH. Statistical significance was assessed using an unpaired two-tailed **t**-test with Welch’s correction. Statistically significant differences are denoted by asterisks (p*≤ 0.05, p**≤ 0.005, ***p ≤ 0.0005). Error bars represent the standard deviation (SD) of the mean, with N = 3 biological replicates.

### Evaluating resistance to transfluthrin using caged mosquitoes in a Peet Grady chamber

The World Health Organization recommends the use of a Peet Grady chamber with caged mosquitoes to assess the efficacy of aerosolized and volatile insecticides^52^. We adapted the methodology described in Silva, Martins et al., 2023^39^ to record differences in susceptibility to transfluthrin between a susceptible (Kisumu) and two resistant (Kisumu-995F and Tiassalé) *An. gambiae* strains. A clear dose-response relationship was observed (Figure 2). The Kisumu strain exhibited substantially higher sensitivity than the resistant strains. The estimated LD₅₀ values (in mg/Whatman paper) were 10.09 (95% CI: 8.74–11.96) for Kisumu and 18.51 (95% CI: 14.85–27.55) for Kisumu-995F. The multi-resistant Tiassalé 13 strain showed minimal mortality even at the highest concentration tested.

**Figure 2.**
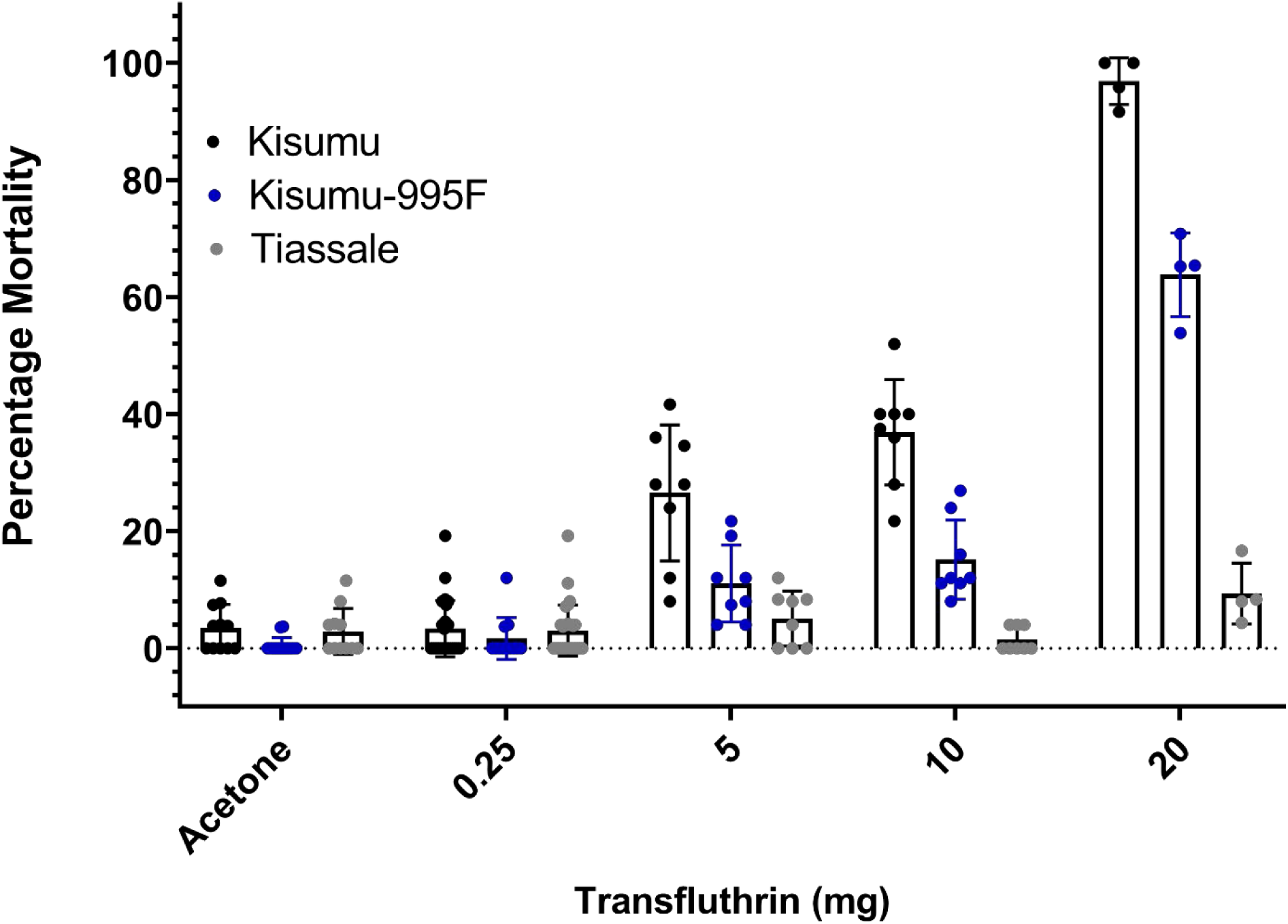
Percentage mortality following exposure to transfluthrin in the Peet Grady Chamber. Values on the x axis represent the total amount of transfluthrin (in mg) applied on the filter paper. Bars indicate standard deviation.

### Video recording mosquitoes’ flight behavior in the presence of transfluthrin reveals differences between pyrethroid resistant and susceptible strains

We video-recorded the flight behavior of caged mosquitoes within the Peet Grady chamber (Supplementary figure 6) in the presence and absence of transfluthrin to assess how exposure impacts their flight behavior and assess differences between pyrethroid resistant (Tiassalé 13 and Kisumu-995F) and susceptible (Kisumu) mosquitoes. Two parameters were evaluated under increasing transfluthrin concentrations: A) the position of mosquitoes in the cage and how it changes over time and B) the total movement of mosquitoes and how it changes over time. Changes in the spatial distribution of mosquitoes in the cage (from the upper part to the floor) may indicate the emergence of intoxication effects of transfluthrin, while an increase in total movement may indicate irritation.

During video analysis each cage half (the left and right sides contain different strains) was split in two compartments: the first (upper) comprised the ceiling, back, and external side wall of the cage, with the second (floor) region comprising the remaining part (Supplementary Figure 6). The mean proportion of mosquitoes occupying each of the two compartments was recorded over time (Figure 3). In the acetone control, more than 75% of mosquitoes in all three strains were located in the upper part of the cage for the whole 60 min of recording. Thus, any increase in the occupancy of the floor is indicative of an impaired flight behavior, due to the insecticide. At the 0.25 mg transfluthrin dose a slight, but steady increase was observed in the number of Kisumu mosquitoes occupying the floor, a small increase in the floor occupancy was observed for Kisumu-995F at the very end of the assay, while the position of Tiassalé 13 remained unchanged. At the higher 5 mg dose a cross over between the blue line, representing the upper part occupancy, and the red line, representing the floor occupancy, was observed for Kisumu at approximately 10 min of exposure (timepoint when 50% of the mosquitoes are on the floor). After approximately 30 min all Kisumu mosquitoes were on the floor, and remained at that position until the end of the 60 min exposure. In the case of Kisumu-995F the two lines converged, at the 5 mg dose, but a cross-over point was only observed at the higher 10 mg dose (after approximately 45 min of exposure). The Tiassalé strain showed only a slight convergence of the lines mainly at the very end of the 60 min at the 5 and 10 mg, but a cross over point was not observed (Figure 3).

**Figure 3.**
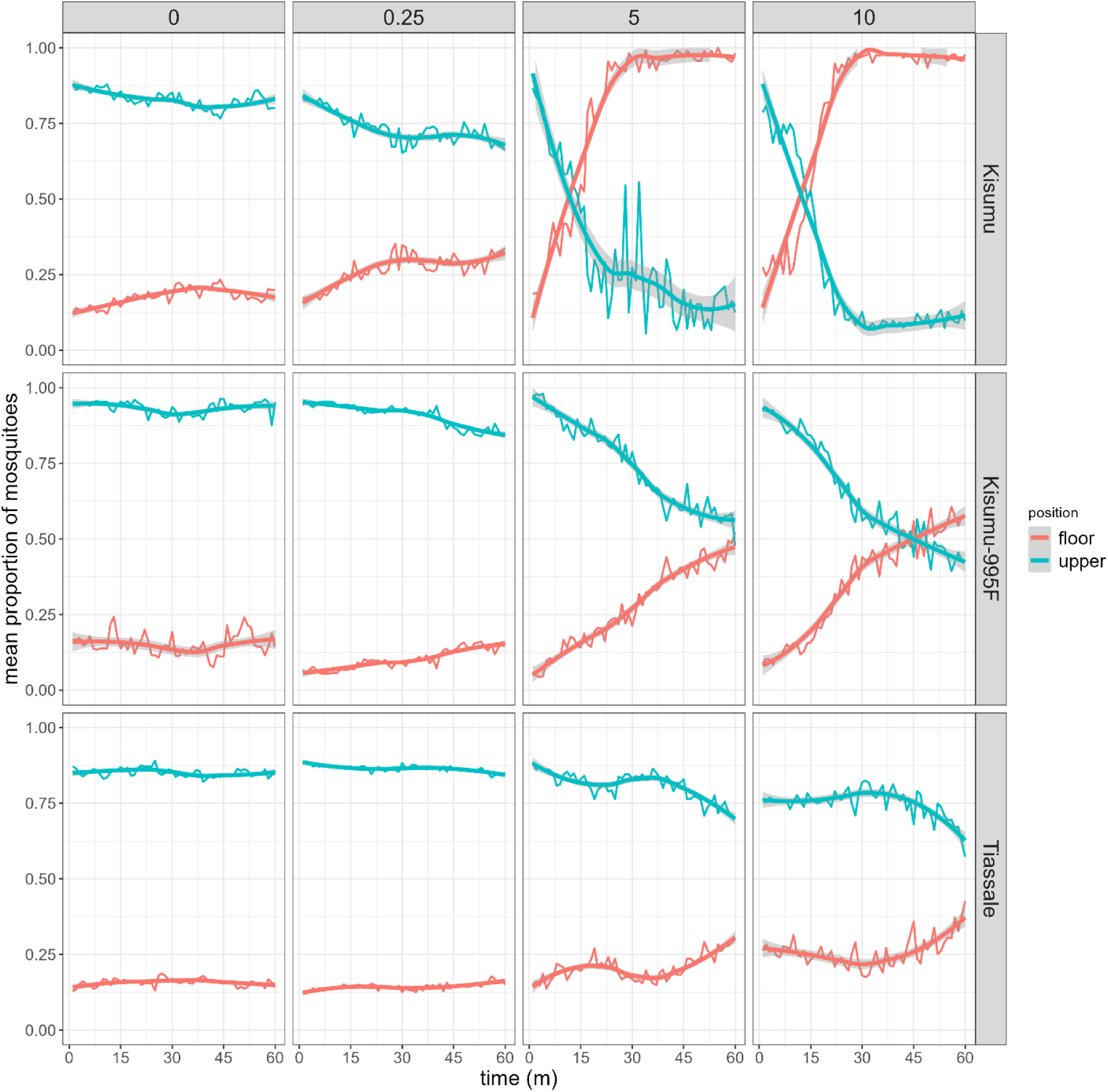
Tracking mosquitoes’ position in the cage during transfluthrin exposure, as a proxy for impaired flight ability. Machine learning analysis of video recordings (at 1 min intervals) to visualize the proportion of mosquitoes occupying the upper part of the cage (blue line) and the floor of the cage (red line) fitted using Loess smoothing method. Data were recorded during 60 min of exposure to different concentrations of transfluthrin (0-acetone control, 0.25 mg, 5mg and 10mg) in the Kisumu, Kisumu-995F and Tiassalé strains.

In addition to tracking mosquitoes’ position, we recorded their movement activity at 5 s intervals to estimate total movement during exposure, serving as an indicator of insecticide-induced movement activation (irritancy) (Figure 4A). During video analysis, each cage was divided into the upper and floor regions, as previously described, and movement was measured both in the whole cage and separately within each region. In the acetone control, Kisumu mosquitoes showed a slightly higher basal level of activity compared to the Kisumu-995F and Tiassalé 13 mosquitoes. At 0.25 mg dose the activity of Kisumu increased and showed a clear difference to the resistant lines, whose activity remained at the basal levels. At the higher 5 and 10 mg doses, the Kisumu line showed a characteristic burst in activity in the first minutes of exposure, which peaked approximately at 10 min and then dropped (more quickly and steeply in the upper compartment), due to the appearance of transfluthrin’s intoxication effects. The two resistant strains also showed a significant increase in movement in both compartments at the higher doses, which increased more steeply in the Kisumu-995F compared to Tiassalé (Figure 4A). We also quantified mosquitoes’ movement in the whole cage. For the higher doses of 5 and 10 mg we quantified the movement only within the first 10 min, as after that time point intoxication effects of transfluthrin become apparent and the movement drops in the susceptible strain due to knock down. For the 0.25mg dose quantification is provided also for the whole 60 min (Figure 4C). The susceptible strain showed significantly higher movement in all doses compared to the resistant lines, as shown in Figures 4B,4C and Supplementary video (with the exception of the comparison with Tiassalé 13 at 10 minutes in the 0.25 mg dose, which had a p-value of 0.14). In the 5 and 10 mg doses the Kisumu-995F showed higher mean movement compared to Tiassalé 13, which was statistically significant only for the 10 mg. The higher movement of Kisumu-995F compared to Tiassalé 13 in the 0.25 mg dose is likely unrelated to transfluthrin exposure, as its levels of movement are lower compared to acetone at that dose and remain unchanged during the 60 min exposure.

**Figure 4.**
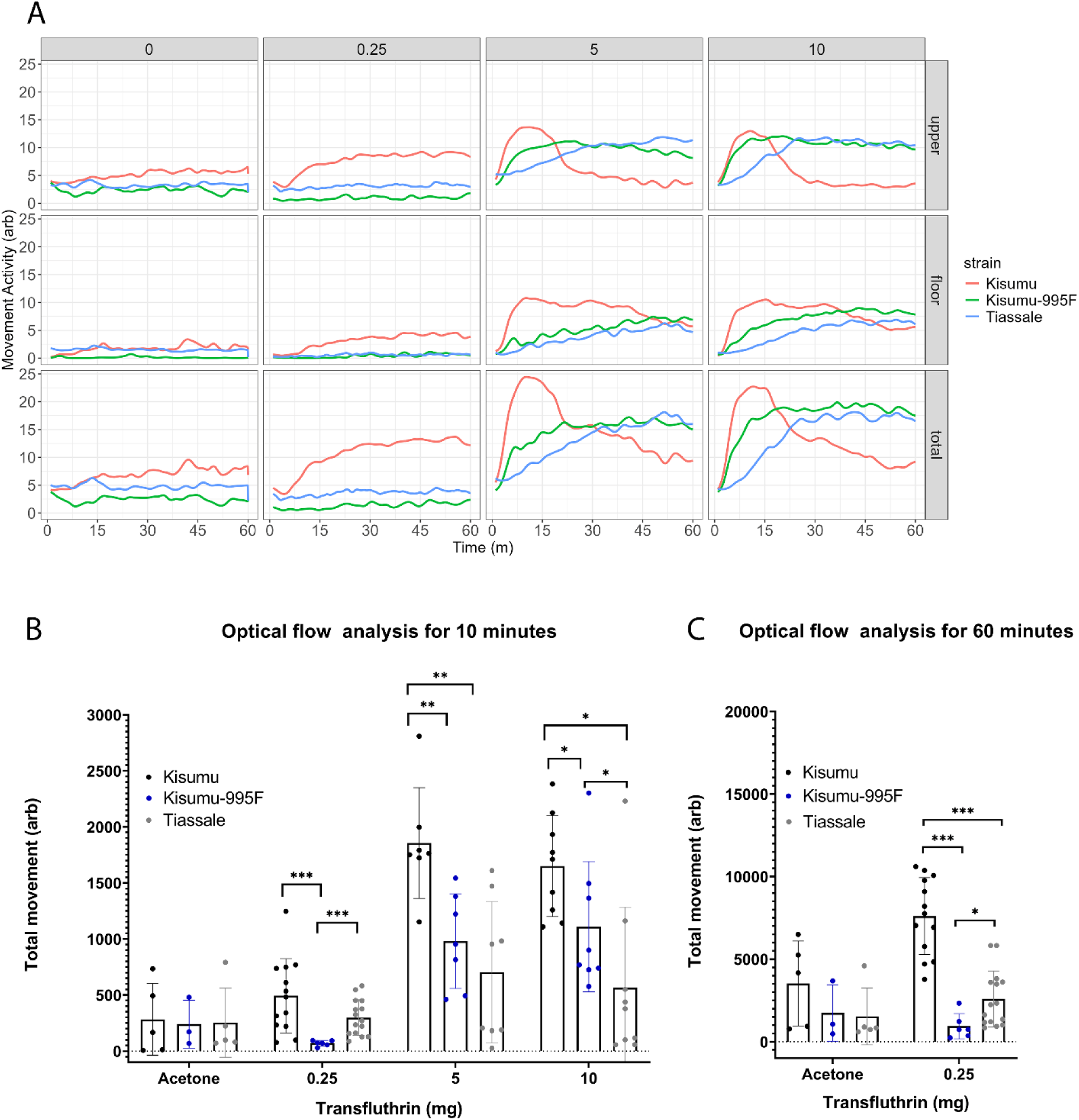
Quantification of mosquitoes’ flight activity during transfluthrin exposure, as a proxy for irritancy. Optical flow method to measure movement activity of Kisumu, Kisumu-995F and Tiassalé 13 when exposed to transfluthrin. A) Movement activity (arbitrary units) measured in 5s intervals, in the upper region (first row), floor region (middle row) and the whole half-cage (last row), during 60 min exposure to varying doses of transfluthrin (0-acetone control, 0.25, 5 and 10mg). B) Mean total movement of mosquitoes (in the whole cage) for the first 10 min of exposure to varying doses of transfluthrin and acetone control. C) Mean total movement of mosquitoes for 60 min of exposure to the sub-lethal 0.25 mg transfluthrin dose and acetone control. For B and C statistical significance was tested using a Mann Whitney U test. Asterisks indicate comparisons with statistically significant difference (*pvalue<=0.05, **pvalue<=0.005,*** pvalue<=0.0005). Error bars represent standard deviations (N=at least 3 replicates of approximately 25 mosquitoes each).

### Resistant and susceptible lines show differences in their blood feeding propensity post transfluthrin exposure

The effect of transfluthrin on female mosquitoes’ blood feeding was assessed immediately (within 1 hour) (Figure 5A) and 24 hours post-exposure (Figure 5B). The Kisumu susceptible mosquitoes showed reduced feeding after exposure to the sub-lethal 0.25 mg dose, which normalized after 24 hours. Resistant strains showed no change at this dose. Kisumu females that were knocked down by higher doses (5 and 10 mg), but recovered within 24 hours, exhibited reduced feeding compared to controls (acetone) and Kisumu exposed to 0.25 mg. Kisumu-995F mosquitoes showed a reduction in feeding immediately after exposure to 5 and 10 mg, which returned to normal after 24 hours. Tiassalé 13 mosquitoes only showed reduced feeding immediately after the 10 mg dose, with no lasting effect.

**Figure 5.**
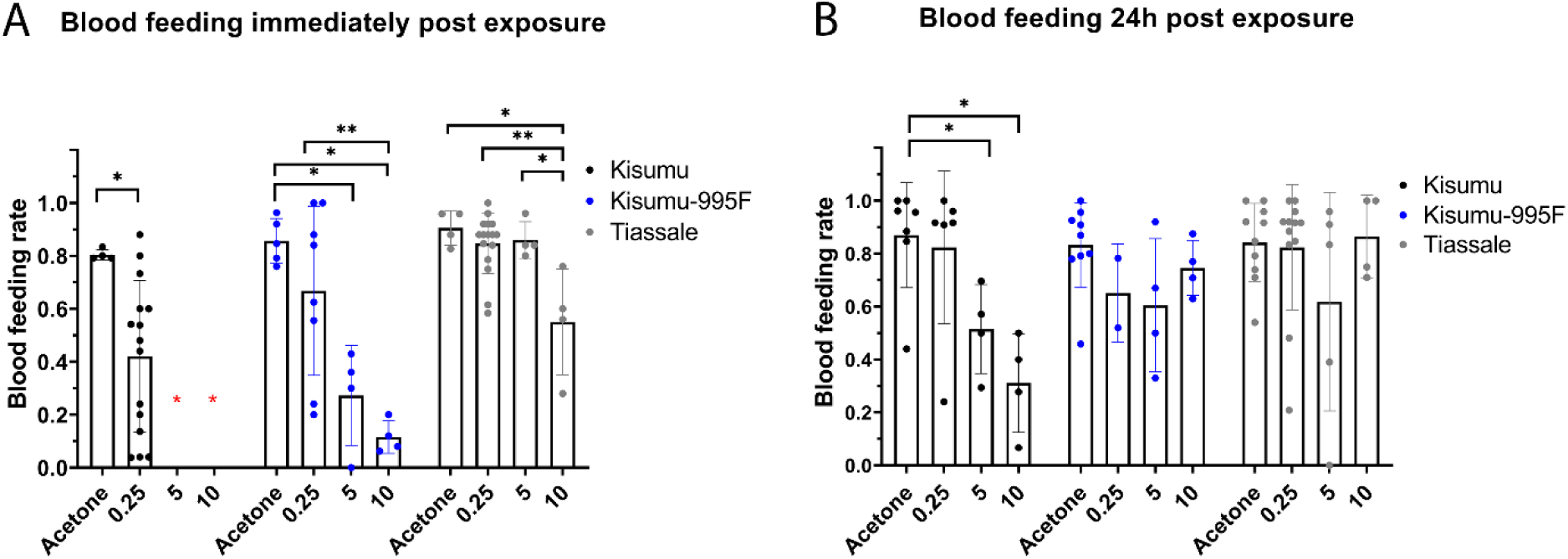
Blood feeding rate upon transfluthrin exposure. The proportion of Kisumu, Kisumu-995F and Tiassalé females that survived transfluthrin exposure and obtained a blood meal: A) 1h after exposure to acetone (control) or varying doses of transfluthrin; B) 24h after exposure to acetone or varying doses of transfluthrin. Different cohorts of mosquitoes were used for A and B. Statistical analysis was performed using a Mann Whitney U test. Asterisks indicate comparisons with statistically significant difference (* pvalue<= 0.05, ** pvalue<= 0.005, *** pvalue<=0.0005). Error bars represent standard deviation (N=at least 3; 25 mosquitoes each). Red asterisks for Kisumu are given for doses that resulted in a high percentage of knock down/death and thus no blood was provided.

### Investigating the role of the *An. gambiae* antennae in transfluthrin detection

To better understand the role of the antennae in the detection of transfluthrin we performed electroantennography experiments and compared the flight behavior of Kisumu mosquitoes with and without ablated antennae in the presence of transfluthrin.

100 and 1000 ppm of transfluthrin (99,11% purity, LGC) diluted in acetone (Pesticide Grade, 99.8% purity, Carlo Erba) were used to test the antennal response of Kisumu mosquitoes. 1-octen-3-ol (>95% purity, LGC) and citronellal (95% purity, Aldrich) were used as positive controls. While a clear response was obtained for the positive controls, transfluthrin elicited no response (Figure 6).

**Figure 6.**
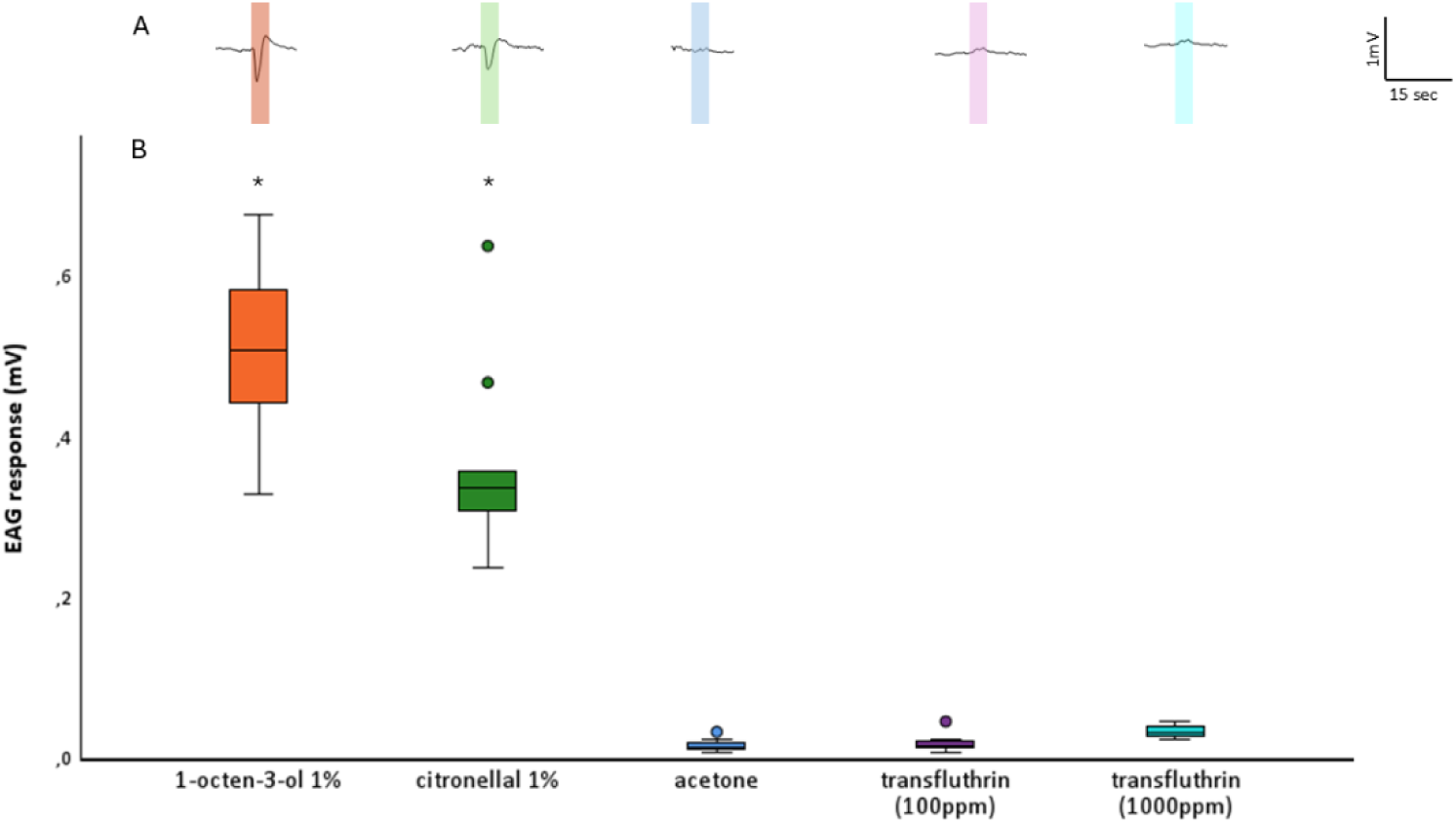
Transfluthrin does not elicit an EAG signal in *Anopheles gambiae*. A. Representative EAG traces for Kisumu antennae exposed to two positive controls (citronellal 1% and 1-octen-3-ol 1%), acetone and two concentrations of transfluthrin (100 and 1000ppm). B. Boxplots of the EAG responses. The bar inside the box represents the median, and the upper and lower parts of the box represent the 25^th^ and 75^th^ percentiles of the data. Circles represent outliers. Asterisks indicate responses that were significantly different from the control (acetone), Kruskal-Wallis Test was performed (F=33.13, df=4, p<0.01) followed by pairwise comparisons where significance values have been adjusted by the Bonferroni correction; 1-octen-3-ol 1% (p<0.001) and citronellal 1% (p=0.049).

We also video recorded the flight behavior of Kisumu mosquitoes with and without ablated antennae in the presence of 5 mg transfluthrin. No significant difference in the total movement was observed between the two groups measured within the first 10 min or over the whole 60 min of exposure (Figure 7A and B) and the flight behavior followed the same trend (Supplementary Figure 7).

**Figure 7.**
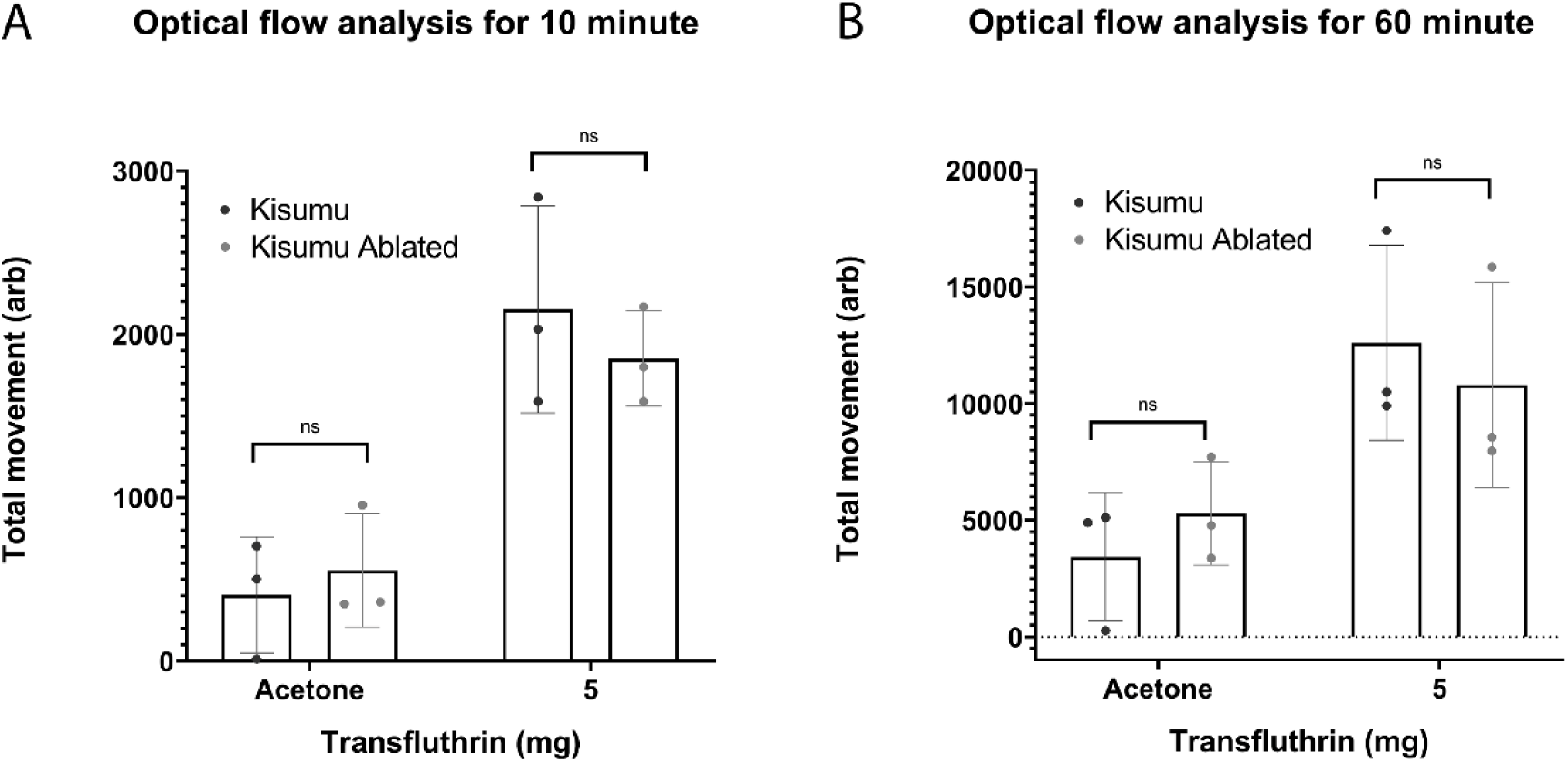
Kisumu mosquitoes with ablated antennae don’t show significant difference in flight activity during transfluthrin exposure. The movement of Kisumu and Kisumu with ablated antennae in the whole cage is quantified during the first 10 min (A) and the whole 60 min (B) of exposure to the acetone control and transfluthrin 5mg. A Mann Whitney U test was performed with p<0.05 significance cutoff, error bars represent SD (N=3; approximately 25 mosquitoes each).

## Discussion

Resistance has been selected to all widely used insecticides in vector control interventions and is increasing both in prevalence and strength^4^. Consequently, any new tool intended for use in public health by targeting the mosquito vector needs to be thoroughly tested for effectiveness in the presence of existing resistance.

Spatial repellents, containing the active ingredient transfluthrin have been prequalified for use in malaria control and are under examination for broader use to control mosquito transmitted diseases^9^. As transfluthrin is a pyrethroid insecticide, the presence and impact of cross resistance on its effectiveness needs to be evaluated. Here we have tested a variety of *Aedes* and *Anopheles* strains resistant to contact pyrethroids for cross resistance to transfluthrin. Overall, our results show that strains resistant to deltamethrin are significantly more likely to be resistant to transfluthrin and those highly resistant to deltamethrin are highly resistant to transfluthrin (Spearman rank correlation ρ=0.76-0.78, p<0.01). This is in line with other studies^53,54^ reporting lower mortality and knock down upon exposure to volatile pyrethroids in strains resistant to contact pyrethroids. However, our results also show that resistance to deltamethrin might not always be a good quantitative predictor of transfluthrin resistance, notable from the results of the *Aedes* strains and the *Anopheles funestus* FuMOZ. It is likely that specific resistance mechanisms or combinations can produce quantitatively different responses to each insecticide. Furthermore, as shown by the contact vs non-contact bioassays, transfluthrin’s route of entry can result in differences in the strength of resistance. This is likely related to the different tissues and resistance mechanisms encountered by the insecticide molecules towards their target.

We also assessed the impact of individual mechanisms on transfluthrin resistance. In line with previous publications suggesting an association of the 995F VGSC mutation with resistance to transfluthrin^19^, our non-contact bioassays using an *An. gambiae* genome edited line carrying this mutation in an otherwise susceptible background, showed that it confers medium levels of resistance (9.3-fold), slightly lower to the previously reported resistance conferred to deltamethrin (14-fold)^35^. The V402L mutation conferred a mild insensitivity (1.7 fold) and like 995F its effect was lower compared to that previously reported for deltamethrin (4 fold)^36^. Contrary to reports suggesting that transfluthrin is not susceptible to P450-mediated detoxification^26,28^, our data indicate significant metabolic vulnerability. Transgenic *An. gambiae* lines over-expressing Cyp6M2 exhibited moderate resistance, while those over-expressing Cyp6P3 show high resistance. *In vitro* metabolism assays demonstrated substantial transfluthrin depletion (65–88%) by *An. gambiae* CYP6M2, CYP6P3, and *An. funestus* CYP6P9a in the presence of cytochrome b5. More modest depletion (<20%) was observed for CYP6P3 and CYP6P9a in the absence of b5, but comparable to the low metabolic activity reported for baculovirus-expressed *An. funestus* CYP6P9a and CYP6P9b lacking b5^28^. This finding highlights the critical role of complete electron-transfer complexes in supporting P450 catalytic efficiency ^55^ and indicates that previously reported inactivity most likely reflected incomplete redox coupling, rather than intrinsic substrate resistance. From a resistance liability perspective, although the fluorinated phenoxybenzyl moiety of transfluthrin is resistant to oxidative attack, metabolism via oxidation of the gem-dimethyl group and formation of the metabolite tetradehydro-2,3-seco-transfluthrin can occur^28^. These alternative metabolic routes may compromise transfluthrin efficacy, and need to be validated through reaction product identification.

Transfluthrin can have multiple entomological outcomes depending on exposure time and concentration, that are important for disrupting mosquito-host interaction. We sought to understand how resistance influences transfluthrin’s effect on mosquito flight behavior and blood feeding. Different experimental set ups have been employed under field and lab conditions to evaluate the repellent effect of transfluthrin^10,56–58^, and there is currently no consensus on optimal methodology, which makes comparison of published results difficult. We used a Peet Grady chamber to video record the flight behavior of caged mosquitoes during transfluthrin exposure. This set up does not permit conclusions to be drawn on the repellent effect of transfluthrin *per se*, as the mosquitoes cannot exit the cage, and we cannot preclude differences in their behavior in the presence of a host and/or other stimuli that would be naturally encountered. However, the assay provides standardized conditions to observe differences between strains in the amount of activity provoked by insecticide exposure, its onset time, peak activity, and the subsequent decline as mosquitoes are knocked-down. The susceptible Kisumu strain showed a higher and quicker activation of movement, compared to the two resistant strains, when exposed to transfluthrin, even at a low dose that did not affect the two resistant lines. This is in line with other studies^20,59^ associating pyrethroid resistance with reduced behavioral avoidance to transfluthrin. However, we also observed differences between the two resistant lines. Kisumu-995F was more affected in the majority of doses tested and showed higher activation compared to the more resistant Tiassalé 13. The activation of movement is likely a sign of neurotoxic irritancy and our data provide evidence that there is a graded behavioral sensitivity that correlates with resistance strength and/or mechanism.

The Peet Grady assay can also be used to test the sensitivity of mosquitoes to the toxic effect of transfluthrin. The effect of transfluthrin exposure is qualitatively different to the knock down response observed in mosquitoes exposed to solid state or contact pyrethroids; instead of rapidly falling to the floor and being unable to stand or fly coherently, the response is slower, with mosquitoes becoming increasingly ‘intoxicated’ and unable to fly high or far but still mobile. This effect can be quantified in real time by tracking the position of mosquitoes in the cage (transition from the upper part to the floor), enabling determination of the dose at which transfluthrin becomes effective in incapacitating exposed mosquitoes. We showed that the effect was dose and time dependent in all strains, occurring at lower doses and more rapidly the more susceptible the strain was. Differences in mortality between strains recorded 24h later were fully consistent with the results of the “deli pot” non-contact bioassay.

We also showed that exposure to transfluthrin can reduce the blood feeding success of mosquitoes immediately post exposure, in a dose dependent manner, with susceptible mosquitoes again being more affected. In most cases the effect did not last beyond 24 h, with the exception of the high doses tested in Kisumu that induced high levels of ‘intoxication’. Thus, transfluthrin concentrations that do not cause mortality, but temporary intoxication can provide prolonged protection by reducing biting. Within small scale laboratory experiments we are able to measure propensity and ability to take a blood meal, which contributes to the blood feeding inhibition induced by transfluthrin, but not the ability to detect and locate a host from a distance for which human landing catch (HLC) or a similar method is needed. Our results are in line with previous studies reporting reduced and delayed blood feeding upon transfluthrin exposure^60^, but importantly reiterate that pyrethroid resistance may at least partially counteract transfluthrin’s effect on blood feeding inhibition.

If and how mosquitoes’ behavioral responses to transfluthrin are underlined by its perception from sensory organs is still largely unknown. Recent studies have suggested that activation of the sodium channel and not of odorant receptors is the primary mechanism of repellency^10^. Our EAG data show that transfluthrin does not elicit a signal in *An. gambiae*, the same as has been reported for *Ae. aegypti*^10^. In addition, ablating the antennae did not result in a significant change in mosquitoes’ activation of movement in the presence of transfluthrin. Thus, our data suggest that mosquitoes’ antennae likely don’t have a primary role in the perception of transfluthrin.

Overall, our data provide evidence that there is a clear risk of reduced efficacy of transfluthrin in the presence of resistance to contact pyrethroids. Further studies on transfluthrin resistance, along with epidemiological measures and modelling efforts are needed to fully assess the impact of transfluthrin resistance on the effectiveness of spatial emanators, especially in terms of reducing disease transmission under different settings, and to evaluate how the risk of reduced efficacy may evolve over time. Molecular markers used to screen field populations for pyrethroid resistance could be used to predict resistance to transfluthrin, but with caution given that the strength of resistance might not be equal and additional mechanisms of resistance could be selected, making different markers relevant for volatile pyrethroids. Research on additional active ingredients with different mode of action to volatile pyrethroids or chemicals that can act as synergists (e.g P450 inhibitors) will benefit the sustainable use of spatial emanators in public health.

## Acknowledgements

This publication is based on research funded by: the Hellenic Foundation for Research and Innovation (H.F.R.I.) under the “3rd Call for H.F.R.I. Research Projects to support Post-Doctoral Researchers” (Project Number: 7406) and the IVCC (through the Gates Foundation [grant number INV-058567]; UK International Development from the UK government [grant number 400499-403]); The findings and conclusions contained within are those of the authors and do not necessarily reflect positions or policies of the Gates Foundation, or the UK government.

We would like to thank David Malone (Senior Programme Officer, Gates Foundation) and Prof. John Pickett for critically reviewing our manuscript.

