## Supplementary Information for "Widely circulating pyrethroid resistance mechanisms reduce the efficacy of transfluthrin and pose a risk for mosquito-borne disease control with spatial emanators"

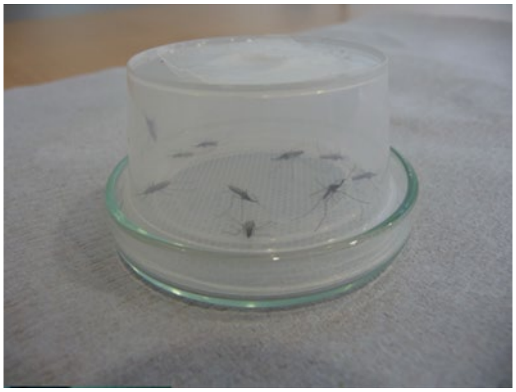

**Supplementary Figure 1**: Deli pot assay with mesh

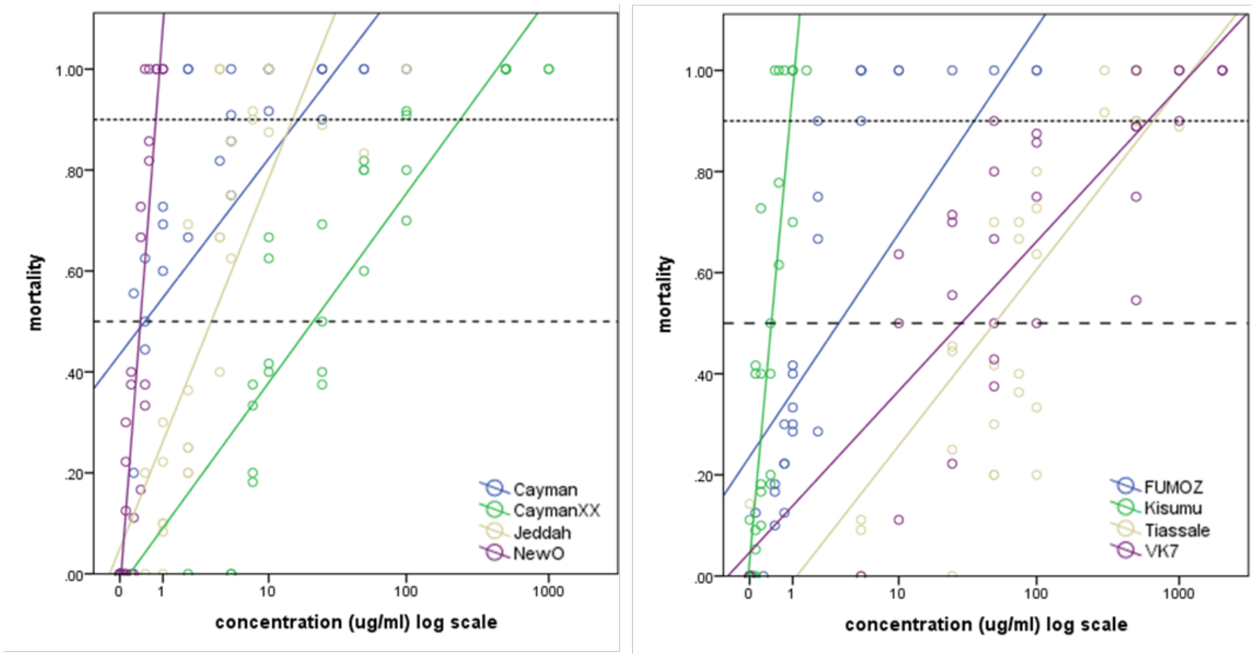


**Supplementary Figure 2**: Transfluthrin *volatile* assay dose-response relationships for each strain (*Aedes* – left panel, *Anopheles* – right panel; the fully insecticide susceptible colonies are NewO and Kisumu). The dashed line shows the 50% mortality threshold and the dotted line the 90% mortality threshold. Note that for *Aedes* Cayman strain, there is negligible difference from the susceptible New Orleans strain in $LC_{50}$ (blue and purple lines almost cross on the 50% threshold) but a much higher resistance ratio if the $LC_{90}$ is considered (large horizontal distance between the blue and purple lines where each crosses the 90% threshold). The plot is indicative of, but may not exactly match the results in the probit regression analysis shown in Table 4.

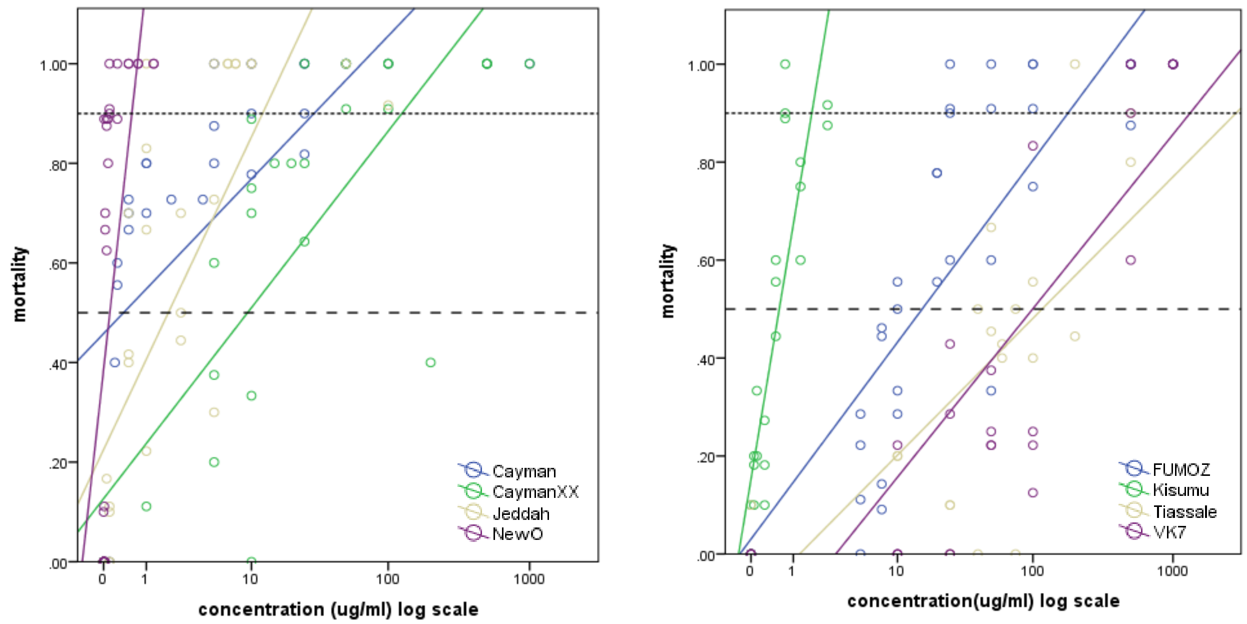


**Supplementary Figure 3**: Deltamethrin tarsal contact assay dose-response relationships for each strain (*Aedes* – left panel, *Anopheles* – right panel; the fully insecticide susceptible colonies are NewO and Kisumu). The dashed line shows the 50% mortality threshold and the dotted line the 90% mortality threshold. Note that for *Aedes* Cayman strain, there is negligible difference from the susceptible New Orleans strain in $LC_{50}$ but a much higher resistance ratio if the $LC_{90}$ is considered. The plot is indicative of but may not exactly match the results in the probit regression analysis shown in Table 3.

resistance ratios in bold are significantly greater than 1 (i.e. significant vs genus-specific susceptible strain).

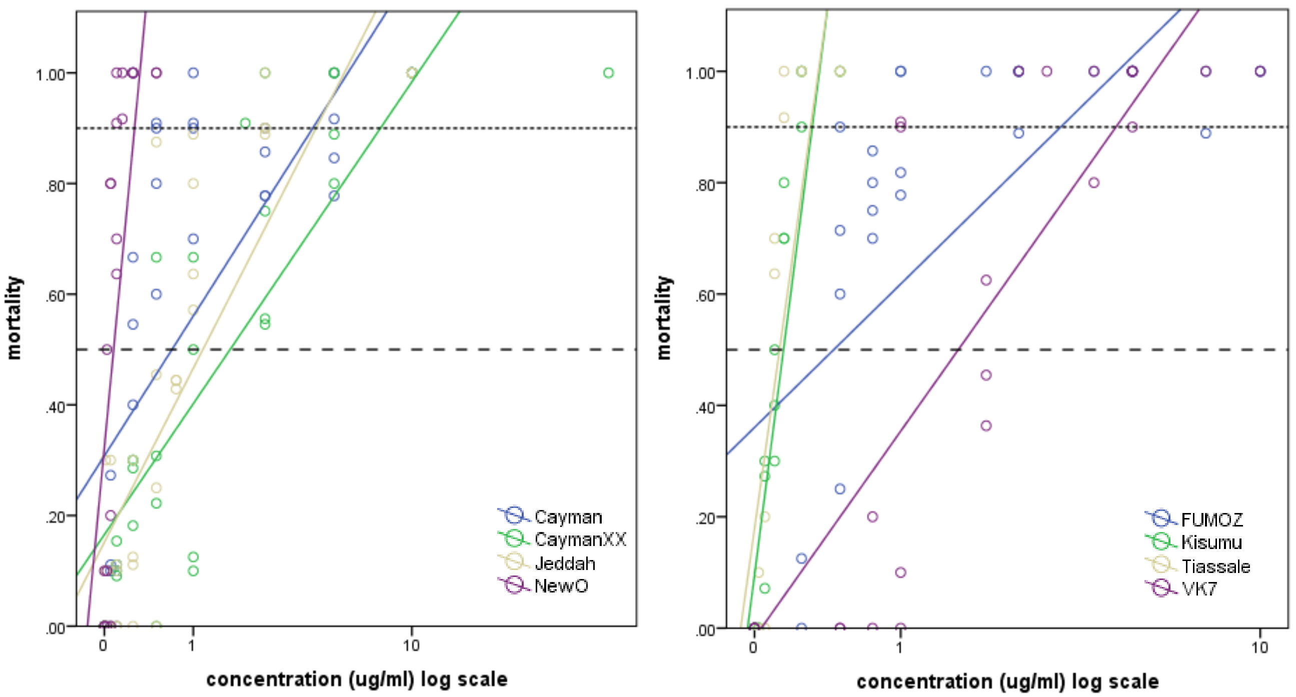


**Supplementary Figure 4:** Transfluthrin tarsal contact (which includes volatile contact) assay dose-response relationships for each strain. Additional explanation is as for Figure 3. Note that in the right hand (*Anopheles*) panel, the data for Tiassale should be ignored as indicated by Table 5 and the strikethrough in the figure legend.

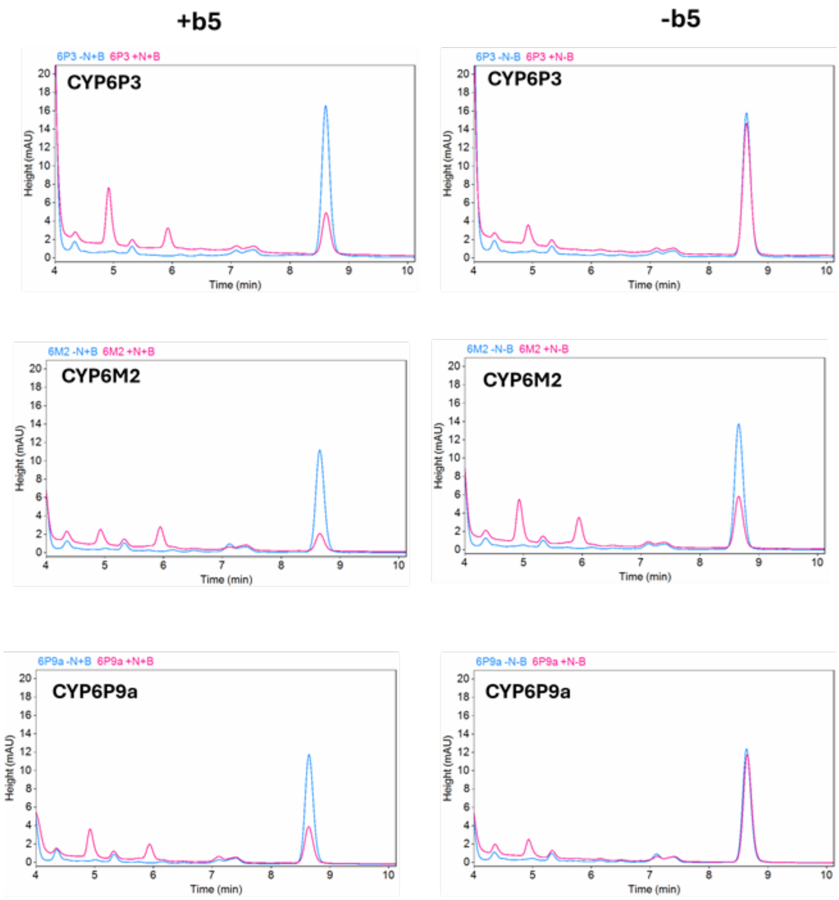


**Supplementary Figure 5**: Representative HPLC chromatograms of transfluthrin metabolism. The overlaid chromatograms represent the results of 2 hour incubations of 100µl reactions containing 0.05Mm P450 plus or minus 0.4µM compound in the presence (pink) and absence (blue) of NADPH.

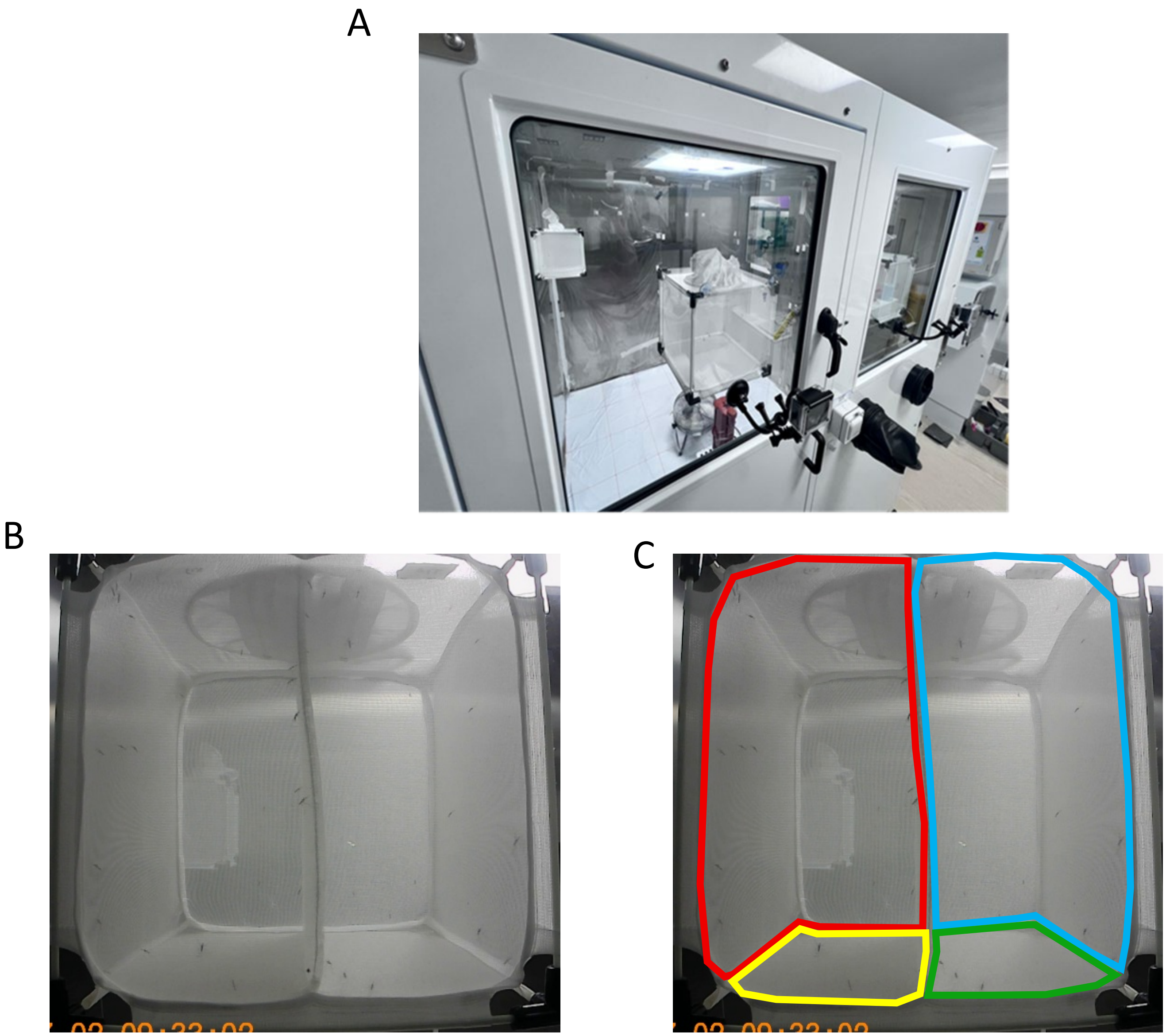

**Supplementary Figure 6**: A) shows an example of a video camera mounted on one of the PG chamber windows. B) shows the actual viewport perspective of the cage from the camera. The area behind each camera was covered by black cotton fabric to minimise external reflections from the outer room. C) Division of the dual-cage mosquito chamber into regions of interest for video analysis. The left strain is grouped by detections in the red and yellow regions. The right strain is grouped by detections in the cyan and green regions. Movement of mosquitoes in the red and cyan regions is registered as 'upper' and movement of mosquitoes in the yellow and green regions is registered as 'floor'.

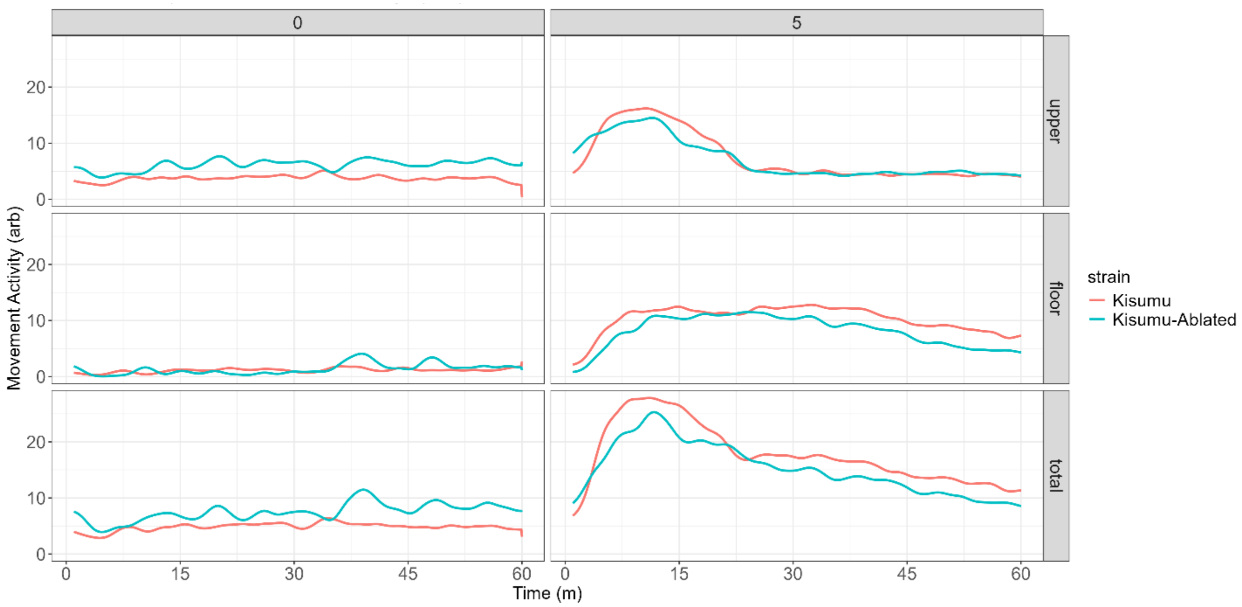


**Supplementary Figure 7:** Optical flow method to measure movement activity of Kisumu, and Kisumu with ablated antennae. Movement activity (arbitrary units) was measured in 5s intervals, in the upper region (first row), floor part of the cage (middle row) and the whole cage (last row), during 60min exposure to acetone (0) and 5 mgr transfluthrin.

| **Species** | **Origin** | **Strain** | **Pyrethroid resistance** | **Resistance mechanisms** |
|---|---|---|---|---|
| *Aedes aegypti* | Grand Cayman | CAYMAN | Resistant | Primarily 3 *kdr* mutations (V410L; V1016I; F1534C) |
| *Aedes aegypti* | Selected from above strain with permethrin | CAYMAN-XX | Strongly resistant | As above + other (likely metabolic resistance) |
| *Aedes aegypti* | Saudi Arabia | JEDDAH | Resistant | Primarily 3 *kdr* mutations (S989P; V1016G; F1534C) + some metabolic resistance (P450s) |
| *Aedes aegypti* | long colonised strain | NEW ORLEANS | Susceptible | None |
| *Anopheles gambiae/ coluzzii* | Côte d' Ivoire | TIASSALE | Strongly resistant | kdr 1014F + metabolic (strong P450; incl. Cyp6P3; Cyp6M2) |
| *Anopheles coluzzii* | Burkina Faso | VK7 | Strongly resistant | *kdr* 1014F + cuticular; some metabolic but not strong P450 |
| *Anopheles gambiae* | Long colonised strain | KISUMU | Susceptible | None |
| *Anopheles funestus* | Mozambique | FUMOZ | Strongly resistant | Cyp6P subfamily P450-based metabolic resistant |

**Supplementary Table 1:** Details of origin, pyrethroid resistance strength and characterized resisatnce mechanisms in strains used for toxicity assays**.**

| **Model** | **Precision** | **Recall** | **mAP50** | **mAP50-95** | **F1 Score** | **Inference Time (ms)** | **Model Size (MB)** |
|---|---|---|---|---|---|---|---|
| YOLOv5n | 0.9940 | 0.9936 | 0.9944 | 0.8523 | 2.98 | 1.2 | 5.268 |

**Supplementary Table 2**: Output parameters of the trained machine-learning model for mosquito detection and localization. The model used was YOLO[40] (You Only Look Once), version 5n, trained and validated on hand-labelled, randomly selected mosquito images from the PG chamber recordings. **Precision** is the proportion of true positive detections among all positive detections made by the model (how many of the detected objects are correctly predicted). **Recall** is the proportion of true positive detections out of all actual objects present (how well the model detects all objects). **mAP50** (mean Average Precision at IoU 0.5) is the average precision across all classes with an Intersection over Union (IoU) threshold of 0.5, where detected bounding boxes overlapping ground truth by 50% or more are considered correct. **mAP50-95** is the mean average precision over multiple IoU thresholds from 0.5 to 0.95 in steps of 0.05, giving a more comprehensive assessment of detection quality by considering stricter matching criteria. **F1 score** is a metric combining precision and recall into a single measure by calculating their harmonic mean. F1 quantifies a model's accuracy in terms of balancing both false positives and false negatives, providing a more comprehensive evaluation than precision or recall alone.

**Inference Time** is the mean time in ms taken to identify all objects in a single video frame. **Model Size** is the size, in MB, of the entire trained network structure and weights in memory.

**Supplementary video:** representative video recording of mosquitoes' movement during exposure to 5mg transfluthrin. The time of recording is between the 5$^{th}$ and 10$^{th}$ min upon initiation of the exposure assay in the Peet Grady chamber. The footage is shown at 16 fold accelerated speed.
